# MicroAge Mission: Effects of Microgravity and Heat Shock Protein 10 Overexpression on the Proteome of Human Tissue-Engineered Muscle Constructs - Implications for Skeletal Muscle Ageing

**DOI:** 10.64898/2026.04.27.721147

**Authors:** Samantha W. Jones, Megan Hasoon, Kareena Adair, Shahjahan Shigdar, Kay F. Hemmings, James R. Henstock, Philip Brownridge, Christopher McArdle, Gianluca Neri, William Blackler, Georgi Olentsenko, Andrew R Jones, Claire Eyers, Kai Hoettges, Malcolm J. Jackson, Anne McArdle

## Abstract

Age-related loss of skeletal muscle mass and function, or sarcopenia, presents a growing clinical challenge, mirroring the accelerated muscle atrophy seen in microgravity. This study, part of the UK Space Agency’s MicroAge Mission, aimed to investigate microgravity-induced proteomic changes in 3D human skeletal muscle constructs and assess whether mitochondrial Heat Shock Protein 10 (HSP10) overexpression could modulate these responses. Constructs derived from control human AB1167 myoblasts and AB1167 myoblasts that were transduced to overexpress HSP10, were flown to the International Space Station (ISS), with a ground reference experiment (GRE) conducted post-flight. Proteomic analysis using mass spectrometry and bioinformatics revealed significant alterations in metabolic, structural, and mitochondrial protein profiles after microgravity exposure.

Microgravity caused downregulation of key proteins involved in energy metabolism, stress responses and structural integrity, while upregulating catabolic and apoptotic enzymes. Many of these modifications parallel previously reported changes in protein composition of muscle with ageing on earth. Overexpression of HSP10 attenuated the effects of microgravity, with fewer proteins showing significant changes and reduced disruption to mitochondrial and cytoskeletal components. Pathway analysis indicated that HSP10 overexpression preserved mitochondrial protein expression, particularly in the matrix, and promoted mitochondrial gene expression and translation under microgravity conditions.

Notably, 284 proteins altered by microgravity in unmodified muscle constructs remained stable in HSP10-overexpressing constructs, suggesting a protective effect. MitoCarta 3.0 analysis confirmed that HSP10 expression modulated protein responses at the mitochondrial level, mitigating declines in bioenergetic proteins that are typically associated with microgravity. Collectively, the findings demonstrate that microgravity induces extensive proteomic remodelling in human muscle, which is partially offset by HSP10 overexpression. These results offer insights into muscle atrophy in spaceflight and suggest that targeting mitochondrial stress pathways via chaperone modulation may be a viable strategy to combat sarcopenia and disuse-induced muscle loss on Earth and in space.

## Introduction

Demographic changes are occurring globally, resulting in greater numbers of older adults (≥ 65 years) in relatively poor health (1) within our populations. One of the most striking consequences of ageing is the progressive loss of skeletal muscle mass and function, a major contributor to disability and hospitalisation in older people and a significant societal challenge (2). It is reported that individuals will lose approximately 3-8% of their muscle mass per decade after the age of 30, with the rate of decline increasing after the age of 60 (2).

In a similar, but accelerated manner, the spaceflight environment poses a significant challenge to the health of astronauts due to prolonged gravitational unloading. Skeletal muscle is particularly vulnerable to the effects of microgravity and undergoes extensive remodeling across short periods of time when compared with muscle during ageing on earth (3). For example, studies have shown that astronauts can lose muscle mass at a rate of ∼ 0.57% per month during long term spaceflight (4–6). Considering these data, there is considerable interest in the possibility that orbital platforms, such as the International Space Station (ISS), could be used as testbeds to study mechanisms of ‘accelerated muscle ageing’ and to develop intervention strategies to prevent muscle deficits in ageing and during spaceflight (7) .

The adaptive capacity of skeletal muscle in response to physical activity and inactivity is governed, in part, by a group of evolutionarily conserved molecular chaperones called Heat Shock Proteins (HSPs). HSPs are key regulators of protein homeostasis and are known to participate in myogenic transcriptional programmes that are activated by both exercise and muscle injury (8). The HSP content of skeletal muscle alters significantly with age and HSPs in muscle show a blunted ability to mount adaptive responses to endogenous and exogenous stimuli (8–11). Further, genetic mutations in HSP genes have been shown to activate atrophic pathways, leading to the development of myopathies (12).

Within the context of spaceflight, studies have shown decreased mRNA expression for HSPs (HSP27, 70 and 84) in rat slow-twitch muscle following exposure to microgravity, which was proposed to be related to a reduction in both neural and mechanical activity (13)

There is increasing evidence that supports a beneficial role for HSP upregulation in the delaying the progression of muscle ageing (12). For example, life-long overexpression of inducible HSP70 (HSP70i) in the muscle of old transgenic mice prevents specific force deficits that are otherwise observed in muscles of old wild-type mice (11, 14). Further, lifelong overexpression of the mitochondrial chaperone HSP10 in mice reduced age-related loss of maximum tetanic force and muscle cross-sectional area (CSA), potentially due to a reduction in the accumulation of oxidatively damaged proteins within the mitochondria (15). These protective effects of HSP10 overexpression support the hypothesis that mitochondrial dysfunction is involved in the development of age-related muscle deficits (15) and mitochondrial defects have also been claimed to underly many of the negative adaptations to spaceflight (16).

The aims of the current studies were therefore to determine the effects of exposure to microgravity during space flight on the ISS on the proteome of tissue-engineered, 3D human skeletal muscle constructs and to determine whether overexpression of mitochondrial HSP10 would modify the microgravity-induced changes. We hypothesized that HSP10 overexpression would reduce the deleterious effects of spaceflight on skeletal muscle in an analogous manner to previous effects seen in muscles from old mice.

## Methods

Three-dimensional, human skeletal muscle constructs, housed in bespoke hardware designed to provide continuous nutrient and oxygen supply, were launched to the ISS aboard SpaceX Cargo Resupply Mission 24 (CRS-24) in December 2021. This formed part of the UK Space Agency-sponsored MicroAge Mission. Full details of the hardware design and the mission have been published (17) and brief details are provided here to allow full interpretation of the data presented.

### Materials

Human immortalised myoblasts (Lot AB1167: *fascia lata* biopsy, 20 years, male), generated using the MyoLine Platform, were obtained via a Materials Transfer Agreement (MTA) from the Institute of Myology (Paris, France) (17–19). Skeletal muscle cell growth medium kits (phenol red-free) were obtained from PromoCell (Heidelberg, Germany).

High glucose, Dulbecco’s Modified Eagle Medium (DMEM) with GlutaMAX™, Leibovitz L-15 Medium with GlutaMAX™, foetal bovine serum (FBS), horse serum, SYBR Green universal master mix, PageRuler™ and high-capacity cDNA reverse transcription kits were purchased from ThermoFisher Scientific, (Altrincham, UK).

The custom lentiviral overexpression vector for Heat Shock Protein 10 (HSP10-T2A-EGFP) and control vector were designed and packaged by VectorBuilder GmbH. PrimePCR SYBR Green qPCR primers (HSP10 and RSP18) and 4–20 % Mini-PROTEAN® TGX™ Precast Protein Gels were purchased from Bio-Rad (Watford, UK). Rabbit polyclonal antibodies to HSP10 (Ab53106), Green Fluorescent Protein (GFP) (Ab6556) were purchased from Abcam (Cambridge, UK). Rabbit monoclonal antibodies to GAPDH (G9545) were purchased from Sigma Aldrich (Dorset, UK). IRDye® 800CW goat anti-rabbit secondary antibodies were purchased from LI-CORbio (Cambridge, UK).

The flight hardware and associated consumables were designed, manufactured or procured by Kayser Space Ltd (Didcot, UK). Unless stated otherwise, all other chemicals used in this study were obtained from Sigma Aldrich, (Dorset, UK).

### Tissue Culture and Fabrication of 3D Muscle Constructs

Human immortalised myoblasts (AB1167) were routinely maintained in a complete growth medium consisting of PromoCell skeletal muscle basal medium (phenol red-free) supplemented with 20 % (v/v) FBS, 50 μg/mL bovine fetuin, 10 ng/mL epidermal growth factor 1 ng/mL basic fibroblast growth factor, 10 μg/mL recombinant human insulin, 0.4 μg/mL dexamethasone, 10 μg/ mL gentamicin and 2 mM L-glutamine. Cells were incubated in a humidified environment at 37 °C with 5 % (v/v) CO_2_, medium exchanges were performed every 48 hours.

Expanded myogenic cell populations maintained in complete growth medium, were dissociated in trypsin-EDTA to a single cell suspension and encapsulated in a fibrin hydrogel solution (consisting of growth-factor reduced extracellular matrix, fibrinogen and thrombin) before dispensing into bespoke, 3D printed scaffolds, polylactic acid (PLA) coated in sterile 0.2 % (w/v) Pluronic™ F-127) at a density of 2.5×10^5^ cells/construct. Each individual scaffold held three muscle constructs, a full breakdown of the fabrication method has been published in detail in Jones *et al*, 2026 (17).

### Stable Overexpression of Heat Shock Protein 10 (HSP10) by Lentiviral Transduction

To generate a cell line that stably overexpressed HSP10, AB1167 myoblasts were seeded sparsely into 6-well plates (1×10^4^ cells/well) in complete growth media and allowed to adhere overnight (37 °C with 5 % (v/v) CO_2_). A lentiviral vector (**Supplementary Figure S1**) was constructed with an EGFP reporter gene, human HSP10 (HSPE1) target sequence, T2A linker and blasticidin resistance gene inserted into the Open Reading Frame (ORF) site. Transcription was under the control of a human EF1A promotor. The plasmid will be referred to as HSP10-T2A-EGFP. Transgene delivery of HSP10-T2A-EGFP was achieved by lentiviral transduction over a period of 24 hours at a multiplicity of infection (MOI) of 10, 50 and 100, supplemented with 5 µg/mL polybrene. Integration of the viral vector into the host cell genome, ensures consistent transgene expression across cell divisions. The following day, the viral solution was replaced with fresh growth medium, and the cells were incubated overnight before undergoing blasticidin (10 µg/mL) selection for 10 days.

All stock cultures (control [AB1167_CTL] and those overexpressing HSP10 [AB1167_HSP10]) were maintained at <70 % confluence and used between passages 4-8 to prevent visual decline of spontaneous contractions. All cultures used for the flight and ground control experiments had >80 % transduction efficiency as confirmed by quantifying the proportion of HSP10-T2A-EGFP expressing cells using a Zeiss Axio Observer Apotome microscope (excitation/emission wavelengths: 488/513 nm).

### Western Blot Analysis of HSP10 Overexpression

Confirmation of HSP10 overexpression was undertaken as previously (15). In brief, 20 µg of AB1167_CTL and AB1167_HSP10 myoblast and myotube lysates in RIPA buffer underwent electrophoretic separation under reducing conditions using a 20% Mini-PROTEAN® TGX™ protein gel. The protein was transferred onto a nitrocellulose membrane using a semi-dry transfer method before staining with ponceau red to ensure transfer uniformity. Membranes were blocked in 5 % (w/v) non-fat milk in 1X Tris-buffered saline with 0.01% (v/v) Tween 20 (TBS-T) for 1 hour at room temperature. Primary antibodies were diluted in 5% (w/v) non-fat milk 1X TBS-T and incubated with membranes overnight at 4 °C (HSP10 1:500, GFP 1:2500 and GAPDH 1:5000). Following washes in TBS-T, membranes were incubated with IRDye® 800 C W goat anti-rabbit IgG secondary antibody (LicorBio), diluted 1:10,000 in 5 % non-fat milk TBS-T for 1 hour at room temperature. Membranes were imaged using the Odyssey CLx Imaging System (LicorBio), densitometric analysis was performed using Image J.

### RT-qPCR Analysis of HSP10 Overexpression

RNA was extracted from AB1167_CTL and AB1167_HSP10 cell lysates using a Qiagen RNeasy Plus Kit, following the manufacturer’s protocol. The quality and quantity of the RNA was assessed using a Nanodrop spectrophotometer. Complementary DNA (cDNA) synthesis was conducted using a high-capacity cDNA reverse transcription kit (Applied Biosystems, ThermoFisher). Amplification of the housekeeping gene and gene of interest was conducted using the ROCHE LC96 light cycler, using a SYBR Green detection system. Primer sequences were proprietary to Bio-Rad^TM^ and not disclosed, product references: PrimePCR™ SYBR® Green Assay: HSPE1, Human and PrimePCR™ SYBR® Green Assay: RPS18, Human.

### Sample Integration into Flight Hardware, Upload and Experimental Configuration

The flight hardware units, manufactured by Kayser Space Ltd (Harwell, UK) are described in Jones et al (17), and designed to interface with the European Space Agency (ESA) incubator, Kubik onboard the ISS. Each flight unit comprised an internal experimental unit (EU) that housed three muscle constructs and autonomously performed medium and fixative flushes following a pre-determined programme.

Sample integration into the hardware at the Kennedy Space Center (KSC) commenced no earlier than 48 hours prior to handover to the launch team. Hardware was launched to the ISS stowed in double cold bags (DCBs) with phase change material (PCM) as previously reported (17). Upload and installation of the hardware into Kubik on the ISS took a cumulative total of 100 hours following the end of sample integration. Upon installation into Kubik each EU was powered, and the pre-programmed experimental timeline was initiated and described in full in (17).

All hardware units immediately executed a medium refresh before the muscle constructs were allowed to acclimate to microgravity on board the ISS for 24 hours. After the acclimation phase, all units performed a second medium refresh. Samples were maintained at 37°C in Kubik for a further 29 hours before the media was displaced with phosphate buffered saline (PBS) and frozen at <-80°C until return from the ISS. Hardware units were returned frozen from the ISS in temperature-conditioned DCBs (-32°C) before being shipped back to the University of Liverpool on dry-ice for sample recovery.

### Ground Reference Experiment

A post-mission Ground Reference Experiment (GRE) was conducted at the University of Liverpool in March 2022. The GRE used the same refurbished hardware and replicated the timings and the temperature profiles of the space-flown experiment on ground (17). Upon completion of the experimental sequence, units were frozen at -80°C before sample recovery.

### Proteomic analysis -Protein Extraction and SP3 Digestion

Muscle constructs were lysed by sonication (Diagenode) in 100 µL lysis buffer (1 % (v/v) SDS, 1 % (v/v) NA-Deoxycholate, 125 mM NaCl, 5 mM EDTA, 100 mM Tris (pH 8), 1X phosSTOP, 1X complete protease inhibitor) before centrifugation (13,000 x g) for 15 minutes at 4°C to remove any insoluble material. The protein concentration of the samples was determined by BCA assay. A total of 7 µg protein for each sample was diluted in 85 µL 25 mM Ambic before a cysteine reduction step was performed by adding 5 µL (11.1 mg/mL) Dithiothreitol (DTT) and incubating the samples at 60 °C for 10 minutes. After cooling, an alkylation step was performed by adding 5 µL (46.6 mg/mL) iodoacetamide (IAM) and incubating for 30 minutes at room temperature under agitation. The IAM was quenched using 4.7 µL (11.1 mg/mL) DTT.

SeraMag™ bead stock was added to each sample (5.26 µL, 50 µg/µL). For protein binding, 420 µL of acetonitrile (80 % (v/v)) was added and incubated for 30 mins at 20 °C. Samples were placed on a magnetic rack to separate the beads from the supernatant. Beads were washed 3 times in 100 % (v/v) acetonitrile before vacuum drying for 10 minutes.

The beads were resuspended in 98 µL Ambic (25 mM) and sonicated for 5 minutes to ensure samples were fully reconstituted. Trypsin stock (2 µL, 0.1 µg/µL) was added to each sample and incubated for 16 hours (37 °C), under agitation. The beads were removed by magnetic separation and the resultant peptide solutions collected in Lobind tubes. Finally, 1 µL 50 % TFA/H_2_O was added to the peptide solutions and incubated for 45 minutes (37 °C) before transferring to 4 °C for a further 30 minutes. Samples were then centrifuged (13,000 x g, 15 minutes) to remove any insoluble material. The cleared peptide solutions were used for LC-MS analysis.

### LC-MS Analysis

The samples (final injection volume 4 µL, 280 ng) were processed using an ultra-high-pressure nano-flow chromatography system (nanoElute, Bruker Daltonics) coupled to a TIMS quadrupole time-of-flight mass spectrometer (timsTOF Pro, Bruker Daltonics) with a modified nano-electrospray ion source (CaptiveSpray, Bruker Daltonics).

The samples were loaded onto trapping columns (Thermo Trap Cartridge, PepMap100, C18, 300 μm x 5 mm) using µL pickup and 4X injection volume plus 2 µL at 217 bar. The samples were resolved on analytical columns (PepSep, twenty-five Series 150 µm, 1.5 µm column) at 35 °C using a gradient of 98 % (A: 0.1 % formic acid)/2 % (B: acetonitrile/0.1 % formic acid) increasing to 65 % A/ 35 % B over 60 minutes at a flow rate of 0.5 μL/min, followed by a washing step at 95% B and column re-equilibration.

Mass spectrometric analysis was performed in dia-PASEF mode. The dia-PASEF method was defined with a m/z range of 475-1000 and a mobility range of 1/ K0 = 0.85-1.3 Vs/cm^2^ using equal ion accumulation and ramp times of 100 ms in the dual TIMS analyser. The collision energy was lowered as a function of increasing ion mobility from 59 eV at 1/K0 = 1.4 Vs/cm^2^ to 20 eV at 1/K0 = 0.6 Vs/cm^2^. The dia-PASEF window scheme consisted of 21 isolation windows of 25 m/z width (cycle time: 0.95 s) with no mass overlap and 1 mobility window.

### Data Quality Check

Raw MS data files were processed using Spectronaut DirectDIA+. Data were searched against three databases: UniProt HomoSapiens, reviewed 12/08/2022 (20,383 entries), UniProt Bovine, 1 protein per gene 05/12/2022 (23,844 entries) and UniProt Mouse 1 protein per gene 05/12/2022 (21,986 entries). The data were searched with the variable modification of methionine oxidation and acetyl (Protein N-term) and a fixed cysteine carbamidomethylation modification and limited to 2 missed cleavages. PSM, peptide and protein group FDRs were set to 0.01 (1%).

### Bioinformatics Analysis

Due to the nature of the muscle construct fabrication process, the samples contained a mix of proteins that aligned to human sequences (derived from the immortalised human muscle cells), and to bovine or mouse sequences (derived from the hydrogel). For proteins with shared sequences, it was not possible to differentiate the protein from the origin species. To overcome this limitation a ‘hydrogel-only’ control sample, containing no human muscle cells, was analysed to highlight proteins contributed by the hydrogel. However, some proteins identified in the hydrogel shared sequences with human proteins found in the samples. In those limited cases, it was not possible to delineate the proportion of a specific protein that was derived from the cells versus the hydrogel within the whole sample.

The proteins were therefore categorised into five groups; *sample* (only human proteins found), *hydrogel* (proteins detected in hydrogel which did not share sequences with human proteins), *mouse/bovine* (proteins which matched only to mouse/bovine but not the hydrogel), *putative_human1* (proteins detected in hydrogel which shared sequences with human), *putative_human2* (human proteins which also matched to mouse/bovine but were not found in the hydrogel). Using these groups, any proteins which were categorised as *hydrogel* and/or *mouse/bovine* were removed, as these were not derived from the human muscle cells. Therefore, all proteins annotated as sample, or *putative human (1 and 2)* remained in the dataset. This resulted in a total of 6060 proteins across the 30 sample set. Distribution of the 30 samples across the experimental groups is shown in **Supplementary Table 1.**

Following exploratory data analysis across the whole dataset using PCA, two samples were removed as they were outliers. Proteins which were missing from more than one sample or were not present in at least two samples within a replicate group (i.e. within one experimental unit – EUID) were removed, resulting in 5067 proteins across 28 samples for the analysis. Data were first corrected using Variance Stabilising Normalisation (VSN). Following normalisation, differential abundance analysis was carried out using the Limma package with FDR multiple test correction. The duplicate correlation function was implemented to correct for the pseudo-replication within EUID groups, with the correlation coefficient being included in the model.

Functional classification and enrichment analysis were carried out using the clusterProflier package. Over-representation analysis (ORA) was carried out for GO terms using the enrichGO function from clusterProfiler. Proteins that were significant in the differential analysis after FDR were used in the ORA and the background consisted of all processed proteins following missing value evaluation and removal.

Kyoto Encyclopedia of Genes and Genomes (KEGG) pathway analysis was carried out using the enrichKEGG function from clusterProfiler. Similar to the ORA, proteins that were significant after FDR were used in the KEGG analysis and the background consisted of all processed proteins following missing value evaluation and removal.

MitoCarta 3.0 (20) containing 1136 human mitochondrial genes, was used to identify the number of mitochondrial proteins present in the total dataset and within each sub-mitochondrial compartment. Following this, the MitoCarta 3.0 list was used to investigate the number of mitochondrial proteins which were differentially abundant across the different comparisons.

All statistical analyses were carried out using R and all figures were created using the ggplot2 package. All code for data analysis and visualisation is available on GitHub (https://github.com/CBFLivUni/MicroAge_Proteomics_paper).

## Results

### Stable Overexpression of HSP10 in Human Immortalised Myoblasts

To generate stable overexpression of HSP10 in human skeletal myoblasts, the lentiviral transduction efficiency of the HSP10-T2A-EGFP plasmid was examined at 24, 144 and 240 hours after blasticidin application and at different multiplicities of infection (MOIs) to select for optimum conditions (10-100; **Supplementary Figure S1 (ii) A-C**). Transduction efficiencies were calculated by counting the proportion of cells that expressed HSP10-T2A-EGFP plotted as a percentage of the total number of cells in the field of view (FOV), The absence of auto-fluorescence was confirmed in non-transduced AB1167_CTL myoblasts (data not shown).

At 24 hours, a MOI of 10 had a transduction efficiency of 46.3 ± 6.8 % which increased to 85.4 ± 8.1 % after 240 hours in the blasticidin selection media. A MOI of 50 resulted in a higher efficiency at 24 hours (67.7 ± 3.3 %), increasing to 95 ± 2.3 % after 240 hours under selection. The greatest starting efficiency was yielded from a MOI of 100 (83.3 ± 19.7) which reached 92.4 ± 3.4 % after 144 hours, this effect was also clearly seen following widefield imaging of the EGFP plasmid in cells (**Supplementary Figure S1 (ii) D**). This condition was selected to produce the stably transduced, mutant cell line (AB1167_HSP10).

To further characterise the cell line, whole cell lysates were subject to analysis via western blot (**Supplementary Figure S1 (ii) E**). Results showed the presence of the EGFP reporter in the AB1167_HSP10 cells as well as a greater abundance (210 % increase) of the HSP10 protein compared with the AB1167_CTL. Gene expression data further confirmed the upregulation of HSP10 by 41.1 ± 21.8-fold in the AB1167_HSP10 line compared with control (**Supplementary Figure S1 (ii) F**).

Independent measurements of the impedance of the muscle constructs obtained prior to freezing on the ISS indicated that the constructs retained viability throughout the mission (protocol reported in full in (17)) and frozen samples recovered from the hardware contained the predicted protein content that was sufficient for proteomics analysis. The protein content of recovered samples is provided in **Supplementary Table ST1**.

### Change in the Proteome of AB1167_CTL Constructs Exposed to Microgravity Compared with the Ground Reference Experiment (GRE)

Comparison of the proteome of AB1167_CTL constructs exposed to microgravity in the MicroAge mission compared with those treated in an equivalent manner in the GRE indicated significant changes in the relative content of multiple proteins detected, with 216 proteins having a significantly higher content in microgravity and 229 having significantly lower content. The remaining 4622 proteins showed no significant changes in content between the two conditions. This is shown graphically as a volcano plot in **Figure 1A** and the top 20 proteins showing changed content are shown as a heat map in **Figure 1B**.

**Figure 1.**
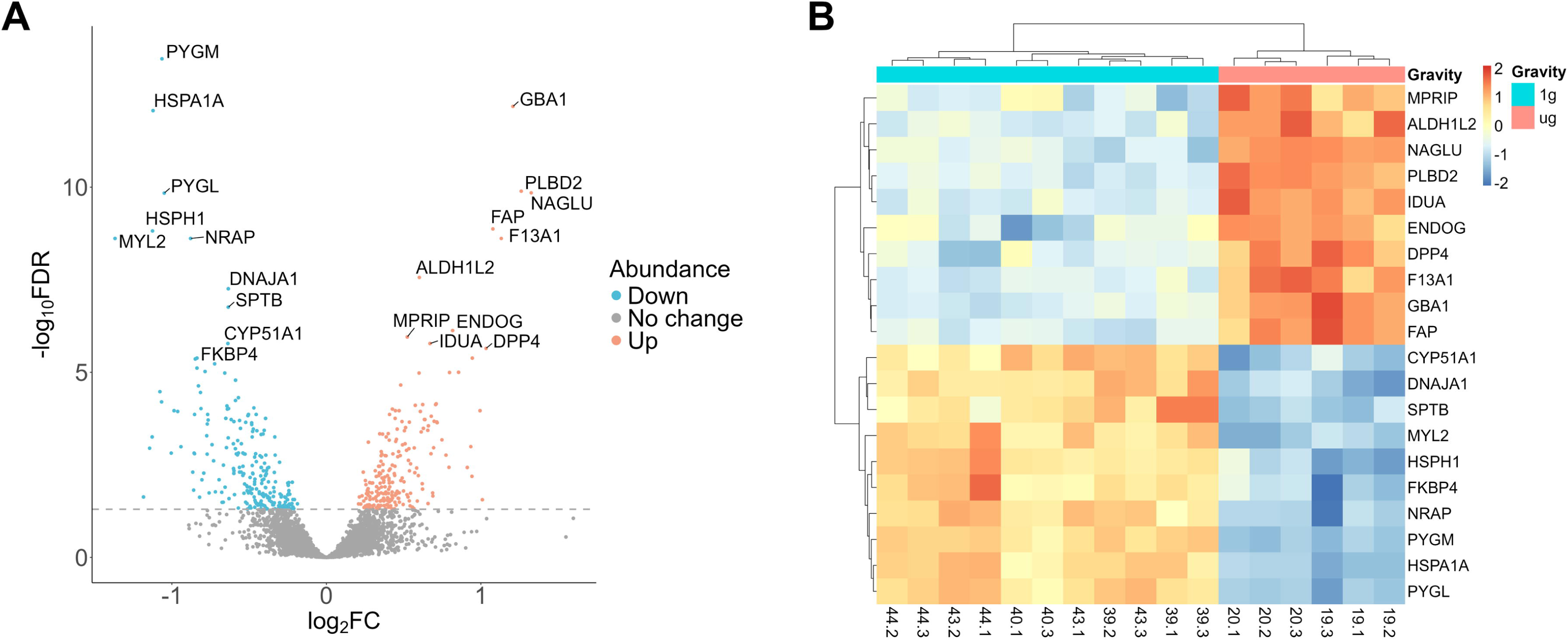
Effect of microgravity on the proteome of AB1167 muscle constructs. A comparison of exposure to microgravity on AB1167 constructs with the protein changes shown as a volcano plot (A) and a heat map of the top 20 proteins modified (B)

The top 10 proteins identified as being decreased in response to microgravity included three HSPs; inducible HSP70 (HSP70i), HSP105 (HSPH1) and HSP40 (DNAJA1), additionally FK506-binding protein 4 (FKBP4), which interacts directly with HSP90. Further, the glycogen phosphorylases PYGM and PYGL, which are essential for gluconeogenesis, were similarly decreased. Structural muscle proteins such as nebulin-related anchoring protein (NRAP) and myosin light chain 2 (MYL2) exhibited reduced abundance in microgravity.

In contrast, proteins increased by exposure to microgravity were predominantly associated with catabolism of macromolecules, including mitochondrial aldehyde dehydrogenase (ALDH1L2), lysosomal N-acetyl-alpha-glucosaminidase (NAGLU) alpha-L-iduronidase (IDUA) and dipeptidyl peptidase (FAP). Enzymes associated with apoptotic cell death and mitochondrial DNA maintenance e.g. endonuclease G (ENDOG) were also increased.

An analysis of the enriched pathways through GO terms and KEGG pathways (**Figure 2 A & B**), confirm modification of multiple metabolic pathway proteins by exposure to microgravity. The output of these pathway analyses shown in **Figure 2 A & B** are non-directional, but an additional comparison of the directional change for the abundance of the specific proteins that contribute to the GO and KEGG pathways allows a determination of whether the changes occur predominantly in one direction. Thus, the identification of significant changes in “cellular lipid catabolic process”, “carbohydrate derivative catabolic process” and “glycosaminoglycan catabolic process” all reflect an increase in the relevant protein contents contributing to increased potential for lipid, carbohydrate and glycosaminoglycan catabolism. In contrast, the identification of changes in “carbohydrate metabolic processes”, “small molecule catabolic process” and “carboxylic acid catabolic process” as major pathways (**Figure 2A**) reflects both relative increases and decreases in the content of relevant proteins. KEGG pathway analysis confirmed an increased capacity of fatty acid and glycan degradation pathways and also identified changes in the content of lysosome-related proteins as a major effect (**Figure 2A**).

**Figure 2.**
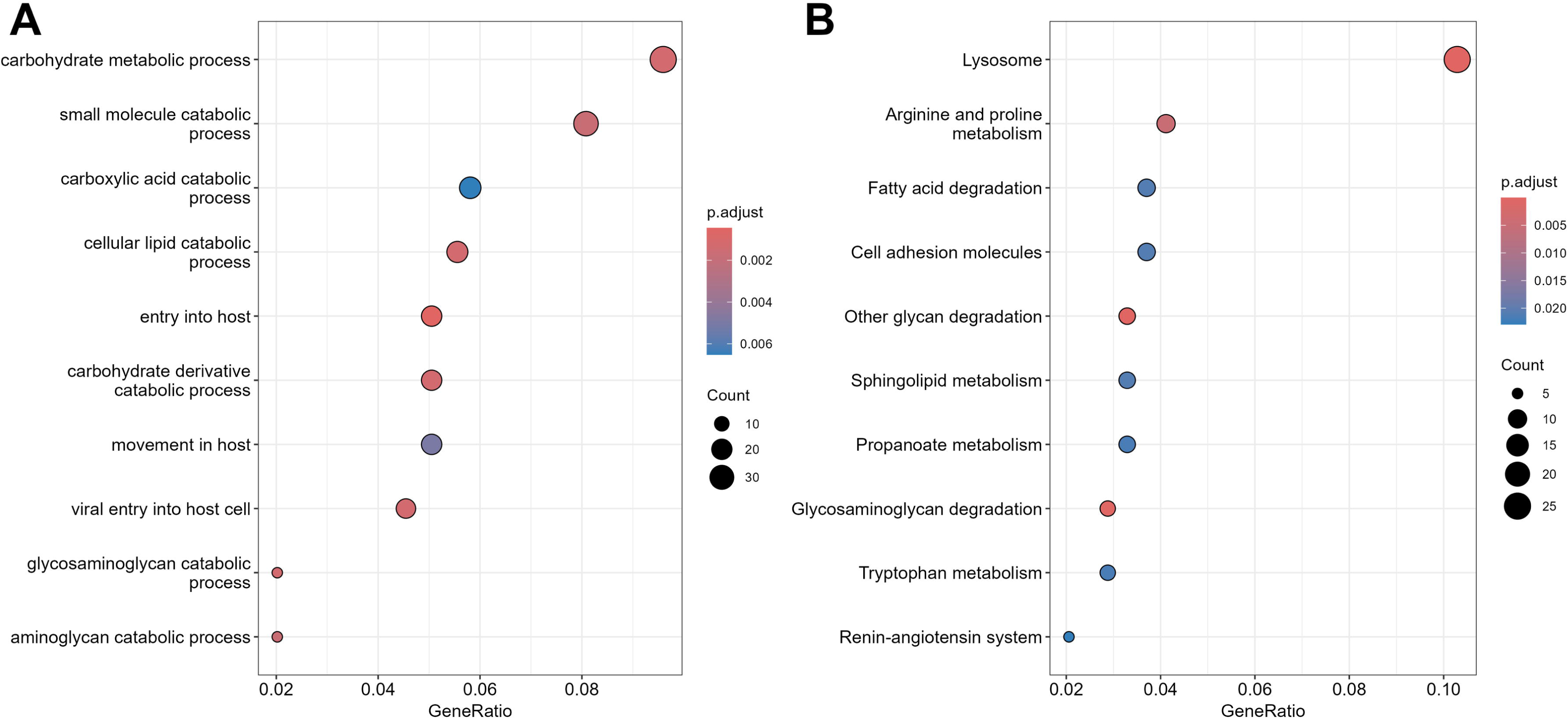
Pathways analysis of the proteome changes caused by caused the effect of microgravity on the AB1167 muscle constructs. Analysis of the enriched pathways using GO terms (A) and KEGG pathways (B) are shown.

### Effect of HSP10 Overexpression on the Proteome of AB1167_CTL Constructs in the Ground Reference Experiment

The effect of HSP10 overexpression on the proteome of AB1167 constructs in the GRE experiment is shown in **Figures 3 A & B** with 255 proteins showing significantly increased content in the AB1167_HSP10 constructs compared with AB1167_CTL constructs and 291 proteins having significantly decreased content. This is shown graphically as a volcano plot in **Figure 3 A** and the top 20 proteins showing changed content are shown as a heat map in **Figure 3 B**.

**Figure 3.**
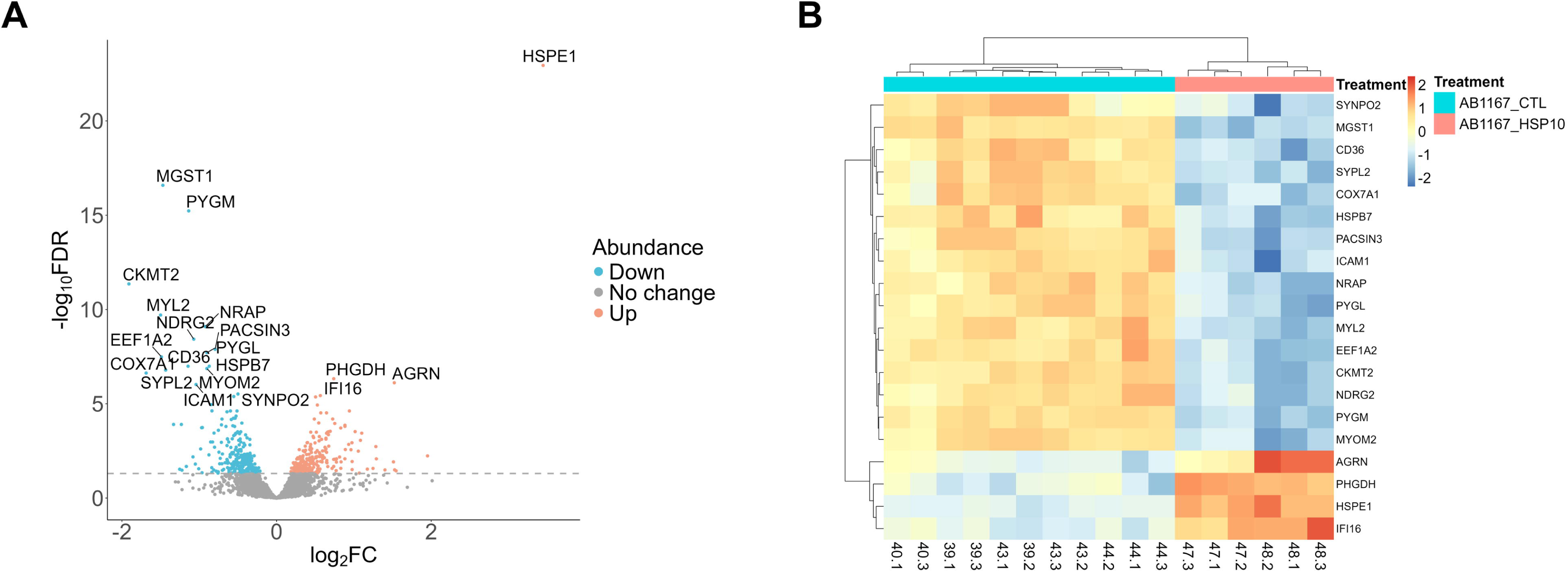
Effect of overexpression of HSP10 on the proteome of AB1167 muscle constructs in GRE. A comparison of differences between proteomes of AB1167_HSP10 constructs and AB1167 constructs in the GRE is shown with the protein changes shown as a volcano plot (A) and a heat map of the top 20 proteins modified (B)

Transgenic upregulation of HSP10 (HSPE1) was confirmed in the proteomic analysis and is clearly seen in **Figure 3A**, with the major effects of HSP10 overexpression appearing to be a decrease in content of multiple metabolic proteins involved in glycolysis (glycogen phosphorylases, PYGL and PYGM), mitochondrial electron transport (COX7A1) and energy supply (CKMT2, mitochondrial creatine kinase and CD36, fatty acid uptake). Further, several structural proteins had reduced abundance, including myosin light chain 2 (MYL2), myomesin-2 (MYOM2) and actin binding proteins such as synaptopodin 2 (SYNPO2), nebulin-related-anchoring protein (NRAP) and Protein kinase C and casein kinase substrate in neurons protein 3 (PACSIN3).

Analysis of the enriched pathways through non-directional GO terms and KEGG pathways **(Figure 4 A & B**), identified both increases and decreases in multiple proteins classified under “system processes” and “morphogenesis”, but the HSP10 overexpression impacted negatively on proteins classified under “muscle structure development”, “energy derivation by oxidation of organic compounds”, “cellular respiration” and changes in “carbohydrate metabolic processes”. The KEGG analysis confirmed these patterns and also highlighted changes in proteins involved in muscle contraction and the citrate cycle.

**Figure 4.**
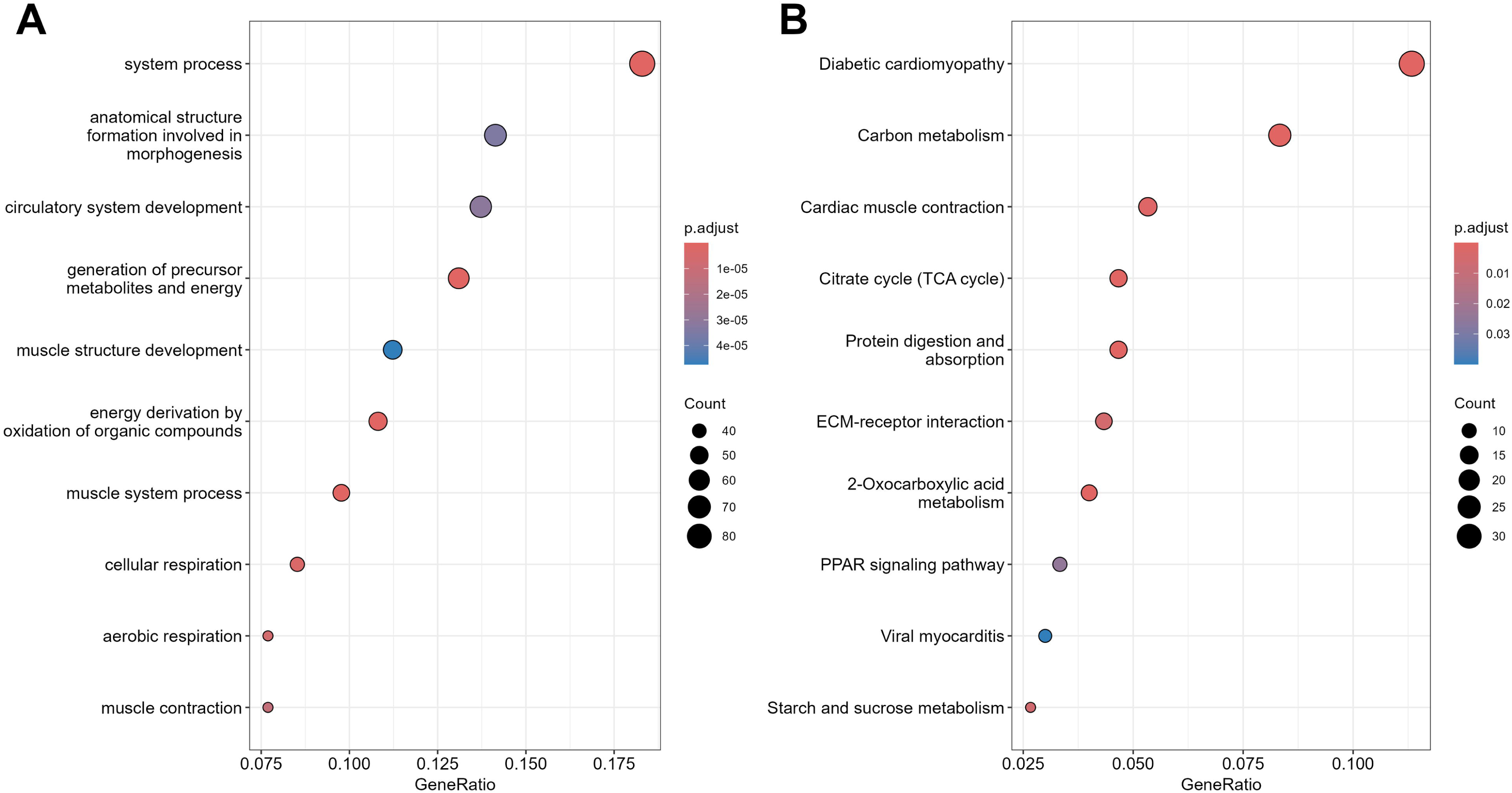
Pathways analysis of the proteome changes caused by the effect of overexpression of HSP10 on AB1167 muscle constructs in GRE. Analysis of the enriched pathways using GO terms (A) and KEGG pathways (B) are shown.

### Change in the Proteome of AB1167_HSP10 Constructs Exposed to Microgravity Compared with the Ground Reference Experiment

The effect of exposure to microgravity was less marked in the HSP10 overexpressing AB1167_HSP10 constructs than was seen in the unmodified AB1167_CTL constructs with only 180 proteins showing a significantly higher content in microgravity and 158 having a significantly lower content (**Figure 5 A & B**). The proteins showing altered abundances on exposure to space flight had considerable overlap with those modified by microgravity in the AB1167_CTL constructs. Again, microgravity exposure resulted in decreased levels of several HSPs, including inducible-HSP70 (HSP70i), HSP105, HSP40 and HSP90. Muscle glycogen phosphorylase (PYGM) was also similarly downregulated. Structural muscle proteins such as myosin heavy chain 2 (MYH2) and myosin binding protein C (MYBPC1) as well as metabolic (carnitine palmitoyl transferase 1A) and redox-related proteins (microsomal glutathione s-transferase 1) were also reduced.

**Figure 5.**
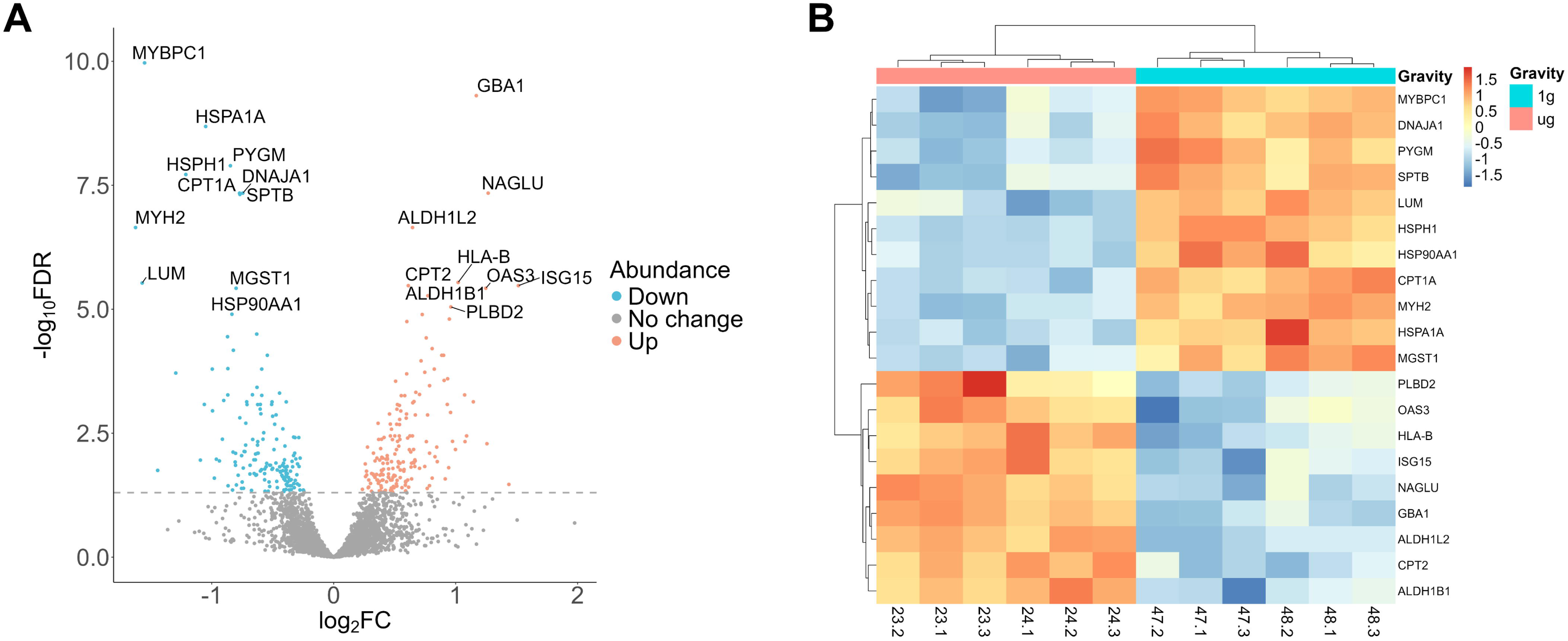
Effect of microgravity on the proteome of AB1167_HSP10 muscle constructs. A comparison of exposure to microgravity on AB1167_HSP10 constructs with the protein changes shown as a volcano plot (A) and a heat map of the top 20 proteins modified (B).

Proteins that were commonly upregulated between the microgravity exposed AB1167_HSP10 and AB1167_CTL constructs included N-acetyl- alpha glucosaminidase (NAGLU), phospholipase B domain containing 2 (PLBD2) and glucosylceramidase (GBA1), again reflecting upregulation of proteins associated with catabolism of macromolecules. The data also showed increased expression of mitochondrial proteins including carnitine palmitoyl transferase 2 (CPT2) and aldehyde dehydrogenase 1L2 and 1B1 in the microgravity-exposed AB1167_HSP10 constructs.

This smaller number of differentially expressed proteins following exposure to microgravity in the AB1167_HSP10 constructs in comparison with the effect seen in AB1167_CTL constructs is reflected in the pathway analyses shown in **Figures 6 A & B**. The GO terms identify increases in multiple proteins involved in mitochondrial gene expression and translation as the major pathways modified by microgravity with decreases in content of proteins associated with muscle tissue development also highlighted. Other pathways highlighted are associated with antigen processing which are likely to reflect the antigenic properties of increased HSP10 expression. Due to the relatively low number of differentially expressed proteins on exposure of the AB1167_HSP10 constructs to microgravity, the KEGG pathways only identified two pathways of which one is associated with energy metabolism.

**Figure 6.**
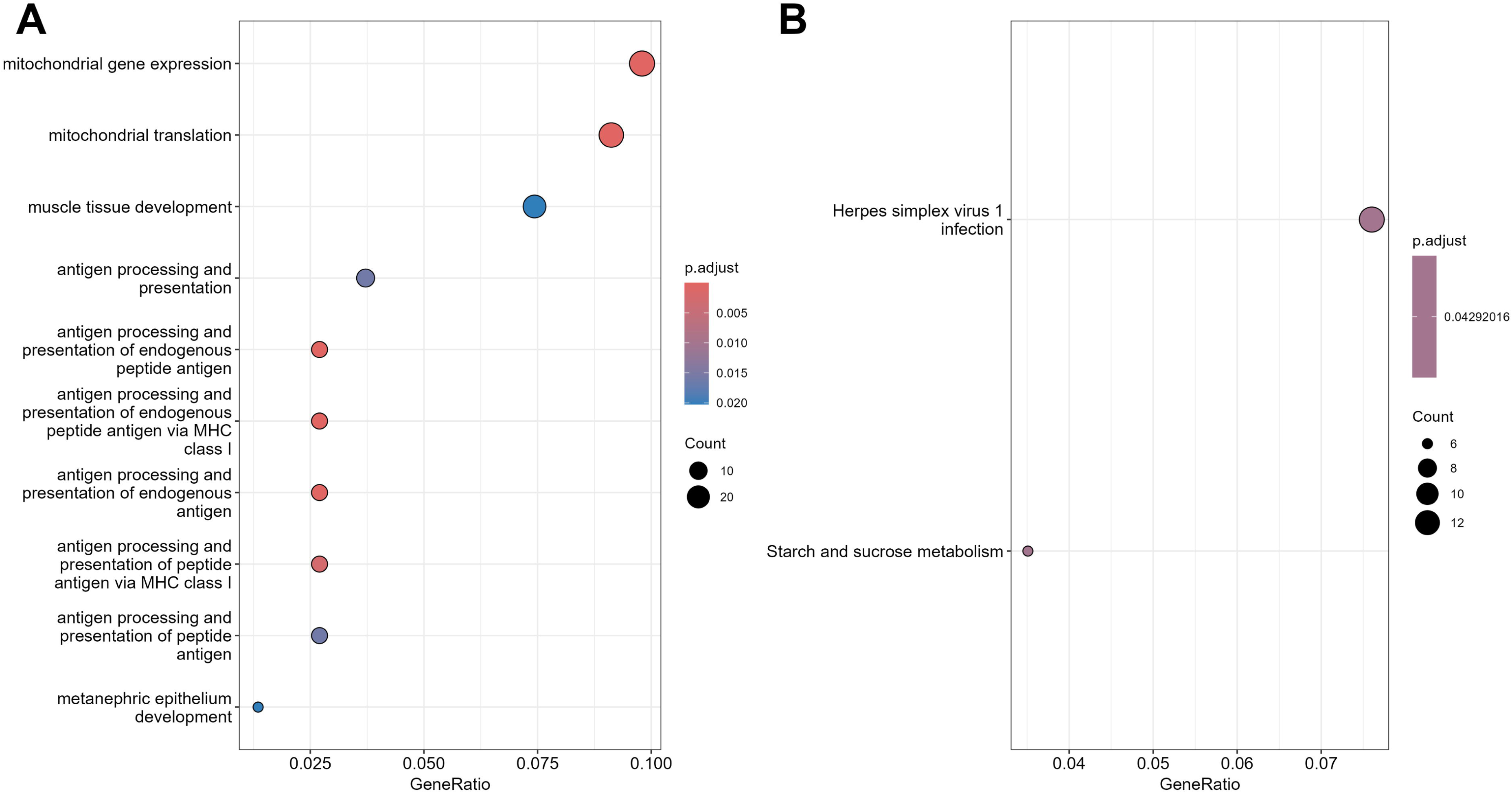
Pathways analysis of the proteome changes due to the effect of microgravity on the AB1167_HSP10 muscle constructs. Analysis of the enriched pathways using GO terms (A) and KEGG pathways (B) are shown.

Despite the commonality of protein changes in response to microgravity in the HSP10 overexpressing constructs compared with those seen in unmodified AB1167_CTL constructs, there were also a large number of proteins (284) differentially expressed following exposure to microgravity in the AB1167 constructs which were not differentially expressed in the AB1167_HSP10 constructs. To address the question of whether this reflects protection by the HSP10 overexpression against the negative effects of microgravity, pathway analysis was undertaken on the 284 unique proteins that were modified on exposure to microgravity in the AB1167_CTL constructs but unchanged in the AB1167_HSP10 constructs on exposure to microgravity. KEGG analyses revealed enrichment in proteins involved in lysosomal degradation, glycan degradation, arginine and proline metabolism and glycosaminoglycan degradation (**Figure 7)**. Three of these pathways (lysosome, glycan degradation and glycosaminoglycan degradation) were identified in **Figure 2B** as being consequences of microgravity exposure in the AB1167_CTL constructs but were not identified in the responses of the HSP10 overexpressing constructs **(Figure 6B**) and thus HSP10 overexpression modified the response of muscle constructs in part by attenuating their metabolic response to microgravity exposure and activation of the lysosomal degradation pathways.

**Figure 7.**
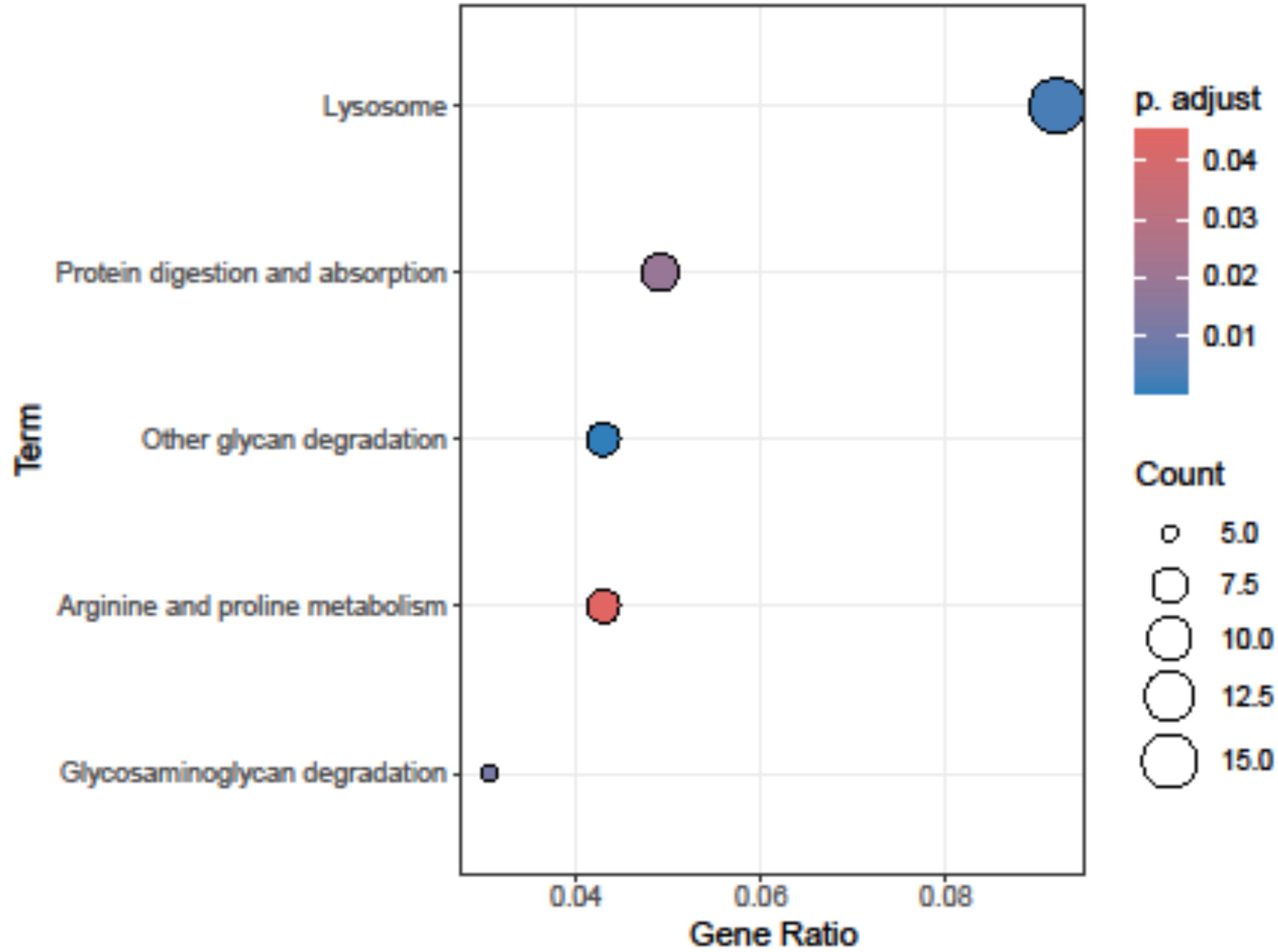
KEG pathway analysis of the 284 proteins that were unchanged in AB1167_HSP10 muscle constructs exposed to microgravity compared with the AB1167 constructs

#### Overlap between protein changes in the various groups

When the four experimental groups were considered together it was apparent that there was substantial overlap between the proteins changed in each comparison and this is illustrated in the “UpSet” plot (21) shown in **supplementary Figure S2**, This plot is an alternative to visualization using a complex Venn diagram and shows the extent of the interactions in these sets of data.

#### The effect of microgravity and HSP10 overexpression on mitochondrial proteins identified using the “MitoCarta 3.0” database

Overexpression of HSP10 in the AB1167 constructs was undertaken as a proof-of-principle approach to determine whether manipulation of mitochondrial stress resistance would modify the responses of the muscle constructs to exposure to microgravity. The data obtained were therefore additionally analysed using the MitoCarta 3.0 database (20) to annotate and select mitochondrial proteins from the data set. The MitoCarta database contains 1136 proteins and 725 of these were found in the 5067 proteins quantified and included in the dataset (i.e. 14.3% of the dataset; **Figure 8A**). Most of the mitochondrial proteins detected in these samples were located in the matrix and inner membrane of the mitochondria in the muscle constructs (**Figure 8A**), but there was no indication of a selective enrichment of proteins in any sub-cellular mitochondrial location. Initial PCA analysis of the MitoCarta data from the 4 groups (**Figure 8B**) showed clear discrimination between the groups of constructs by both the presence or absence of gravity and presence, or absence of HSP10 overexpression. Thus, both variables had substantial regulatory effect on the contents of mitochondrial proteins in the constructs.

**Figure 8.**
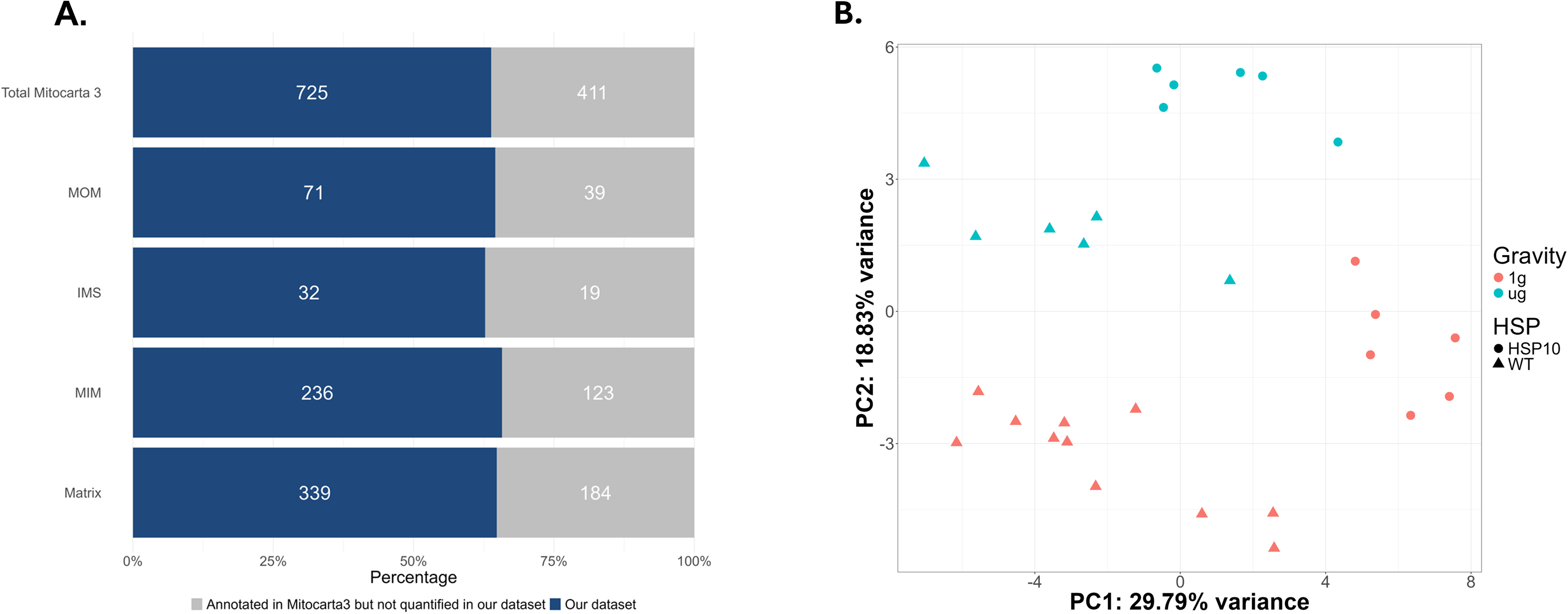
Bar diagram showing a comparison between the number of proteins in the MitoCarta 3.0 database and the number detected in the current data, together with sub-cellular location of the proteins detected (A). PCA of the mitochondrial proteins identified from the Mitocarta 3.0 database that were detected in the 29 samples analysed, showing clear discrimination of the experimental groups (B). Key. MOM: mitochondrial outer membrane; IMS: Inter membrane space; MIM: mitochondrial inner membrane.

Heat maps of the top 20 modified mitochondrial proteins following exposure of AB1167_CTL constructs or AB1167_HSP10 constructs to microgravity are shown in the supplementary data (**Supplementary Figure S3A & B)** and an examination of the number of mitochondrial proteins showing significant changes on exposure of AB1167 constructs to microgravity indicated that of the 445 proteins showing significant changes (reported in **Figure 1**) 78 were mitochondrial proteins (17.5%) (**Table 1**), whereas the number of mitochondrial proteins showing significant changes on exposure of AB1167_HSP10 constructs exposed to microgravity (**Figure 3**) was 77 in the group of 338 significantly changed proteins (22.7%).

**Table 1.** Mitochondrial localisation of proteins found in the *Mitocarta 3.0* database. **Abbreviations.** IMS: Inter-membrane space; MIM: Mitochondrial inner membrane; MOM: Mitochondrial outer membrane

| Comparison | IMS | Matrix | Membrane | MIM | MOM | Unknown | Total number of proteins in *Mitocarta 3.0* |
| --- | --- | --- | --- | --- | --- | --- | --- |
| AB1167 GRE vs μg | 3 | 45 | 3 | 15 | 9 | 3 | 78 |
| AB1167_HSP10 GRE vs μg | 5 | 57 | 1 | 8 | 4 | 2 | 77 |
| AB1167 vs AB1167_HSP10 in GRE | 7 | 57 | 4 | 45 | 10 | 3 | 126 |

Catalase (CAT), carnitine palmitoyl transferase A1 (CPT1A), and microsomal glutathione S-transferase 1 (MGST1) were consistently decreased in response to microgravity, irrespective of HSP10 overexpression (**Supplementary Figure S3A & B**). This indicates that microgravity exerts a deleterious effect on both fatty acid metabolism and antioxidant capacity. However, additional deleterious impacts were seen in the unmodified AB1167_CTL constructs in response to microgravity that were not identified in the AB1167_HSP10 comparison, such as downregulation of HSP60 (HSPD1), carnitine palmitoyl transferase B1 (CPT1B), and mitochondrial creatine kinase.

Conversely, several protein degradation enzymes were consistently increased, including leucine aminopeptidase 3 (LAP3) and alpha-aminoadipic semialdehyde synthase (AASS). Notably, constructs expressing AB1167_HSP10 also exhibited a pronounced increase in ribosomal-associated proteins which were not seen in the control constructs (**Supplementary Figure S3A & B**).

An examination of the effect of HSP10 overexpression in the GRE using MitoCarta showed a significant change in 126 mitochondrial proteins (23.1% of the total number of proteins) The top mitochondrial protein changes in the HSP10 overexpressing constructs in GRE are shown as heat maps in **Supplementary Figures S4.** HSP10 overexpression showed notable effects on mitochondrial metabolism, specifically upregulation of carnitine palmitoyltransferase 1A (CPT1A) and short-chain dehydrogenase/reductase 1 (DHRS1), suggesting enhanced fatty acid oxidation and lipid metabolism. Conversely, several other critical proteins involved in mitochondrial function and bioenergetics were decreased.

The MitoCarta 3.0 database also provided information on the sub-cellular mitochondrial location of the proteins modified by exposure to microgravity and hence a potential indicator of likely sites of mitochondrial remodeling induced by the variable. **Table 1** shows the numbers of proteins in each of these sites for the four comparisons undertaken. While the number of proteins changing was relatively small, it appears that exposure to microgravity primarily caused a substantial change in matrix proteins, whereas HSP10 overexpression affected both matrix and outer mitochondrial membrane proteins in the GRE.

## Discussion

The aims of this study were to determine the effects of exposure to microgravity during space flight on the ISS on the proteome of tissue-engineered, 3D human skeletal muscle constructs and to determine whether overexpression of mitochondrial HSP10 would modify the microgravity-induced changes in an analogous protective manner to previous effects seen in muscles from old mice. Overall, the proteomic data demonstrate that microgravity induces extensive proteomic remodelling in human muscle, which was partially offset by HSP10 overexpression.

Formation of muscle constructs suitable for space flight from the immortalized human muscle myotubes required their packaging in an extracellular matrix derived from a hydrogel (17, 19). Inevitably the hydrogel proteins were also analysed in the protein mass spectrometry and the bioinformatic analysis was developed to remove these confounding proteins. A ‘hydrogel-only’ control sample was analysed to identify which proteins were present and a limited number of proteins were identified in the hydrogel that shared sequences with human proteins detected in the muscle samples. In those limited cases, it was not possible to delineate the proportion of a specific protein that was derived from the cells versus the hydrogel within the whole sample and they were excluded. In practice, any proteins which were categorised as *hydrogel* and/or *mouse/bovine* were removed, as these were not derived from the human muscle cells. This resulted in identification of a total of 5067 human muscle proteins across the 29 sample set which were used in the bioinformatic analysis.

The major changes in the proteome of the AB1167_CTL constructs on exposure to microgravity reflect reduced content of proteins involved in muscle structure and contractions (e.g. NRAP, MYL2) (22, 23) and reduced chaperone (HSP70i, HSP105, HSP40) levels linked to muscle atrophy and reduced stress responses (12). These changes were associated with changes in endonuclease G, mitochondrial nuclease that acts as an effector of apoptosis (24). The pathway analyses highlighted substantial changes in energy metabolism of the muscle constructs with a reduced capacity for glycogen utilization through declines in glycogen phosphorylase content associated with an increased capacity for catabolism of multiple macromolecules and modified lysosomal activity. Our interpretation of this is that the proteome changes reflect a switch in metabolism to maintain available energy supply to the constructs. Changes in the fuel utilization capacity of muscles have been previously reported in proteomic analyses of rodent muscles exposed to space flight (25) and reductions in HSP expression have been reported in multiple models of disuse atrophy (26–28).

Previous studies have also highlighted disruption of mitochondria as a potential cause of energy supply disruption in space flight (16, 29, 30) and the proteomic changes induced by exposure to space flight were therefore also examined using the MitoCarta3.0 database (20) to identify the mitochondrial proteins in the dataset. This analysis indicated a moderate relative enrichment of mitochondria-localised proteins in the spaceflight-induced changes with the proportion of mitochondria-localised proteins increasing to 17.5% from 14.3% in the total dataset. The majority of these proteins being localized to the mitochondrial matrix (Table1). These data are therefore fully in accord with the effects of microgravity affecting the content of proteins involved in energy supply to the muscle myotubes with a particular effect on mitochondria matrix localized proteins.

Exposure to microgravity or space flight have been proposed as accelerated ageing models for skeletal muscle and it is relevant to compare the changes reported here to previous studies of the effects of ageing on human muscle. Clear parallels can be seen in the reduction in the HSP content of skeletal muscle in ageing (8–11) and the changes with microgravity reported here and muscle from ageing individuals also show changes in proteins related to energetics with reduced glycolytic capacity (31, 32). These changes in ageing muscle were associated with reductions in structural proteins, such as myosin light chains which were also seen here. Markedly, exposure to microgravity and ageing are both reported to be associated with disruption of muscle mitochondrial function (16, 31–33)and the data presented here are fully in accord with that conclusion.

Because of disruption of mitochondrial function has been linked to the mechanisms underlying muscle loss following both exposure to space flight and ageing on earth, it was hypothesised that muscle constructs overexpressing the mitochondrial co-chaperone HSP10 would show a reduced response to exposure to microgravity in space flight in an analogous manner to the previously described effects of HSP10 overexpression on age-related declines in mouse skeletal muscle (15).

Mitochondria contain several chaperones, including HSP60, HSP10 and Grp75. HSP60 and HSP10 together form the HSP60/10 chaperonin complex, which is responsible for the folding of proteins transported into the mitochondrial matrix, refolding, and prevention of aggregation of denatured proteins (34). In studies of muscle ageing, mitochondrial HSP10 overexpression was found to prevent the increase in markers of oxidative damage in mitochondria usually seen in older wild type mice (15).

Overexpression of HSP10 in AB1167 constructs in the GRE led to substantial protein changes **(Figure 3).** In a similar pattern to the effects of microgravity exposure on AB1167 constructs, the HSP10 overexpression surprisingly caused a decrease in glycogen phosphorylase and other proteins involved in short term energy supply (e.g. creatine kinase). The GO analysis also indicated a decline in various energy-generation pathways (“energy-derived by oxidation of organic compounds” and “cellular respiration”). These data suggest that the HSP10 overexpression reduced the capacity of the muscle constructs to undertake glycogenolysis and mitochondrial fatty acid metabolism. Several structural muscle proteins (MYL2, MYOM2, NRAP) were also down regulated by the HSP10 overexpression potentially reflecting an effect of the overexpression on proteostasis (35). Analysis using the MitoCarta3.0 database indicated that a high proportion of the significant protein changes were in mitochondria-localised proteins (23.1%) with these proteins being predominantly localised to the mitochondrial matrix (57/126) and mitochondrial inner membrane (45/126). Thus, it appears that overexpression of HSP10 in constructs in the GRE prior to spaceflight induced multiple comparable changes in glycolytic capacity, structural proteins and mitochondrial proteins that were also seen on exposure of control AB1167 constructs to microgravity.

These changes caused by HSP10 overexpression caused a significant modification of the responses of AB1167_HSP10 constructs to microgravity compared with AB1167_CTL constructs **(Figures 5 and 6).** The number of differentially expressed proteins reduced from 445 in the AB1167_CTL constructs to 338 in the AB1167_HSP10 constructs

Some of the changes induced by microgravity in the AB1167_CTL constructs were also seen in the AB1167_HSP10 constructs including the depression of content of the chaperones (HSP70i, HSP105, HSP40 and HSP90), together with evidence for reduced muscle and metabolic proteins (PYGM, MYH2, MYBPC1) reflecting the persistence of microgravity-induced metabolic stress and atrophy. A substantial change in response of the AB1167_HSP10 constructs was however reflected in the GO and KEGG pathway analysis which show increases in mitochondrial gene expression and mitochondrial translation resulting from the microgravity exposure in the HSP10 overexpressing constructs that were not seen for the AB1167_CTL constructs.

Overall, 284 proteins were differentially expressed by exposure to microgravity in the AB1167_CTL constructs that were not differentially expressed in constructs with HSP10 overexpression. Pathway analysis of the 284 proteins which were differentially expressed by exposure to microgravity in the AB1167 constructs but not the HSP10 overexpressing constructs revealed an enrichment in pathways related to lysosomal degradation, glycan biosynthesis and metabolism and amino acid metabolism. The MitoCarta analysis also supports this effect of HSP10 overexpression on mitochondrial proteins with 22.7% of the significantly modified proteins following HSP10 overexpression being localized to the mitochondria, primarily to the mitochondrial matrix (57/77).

A comparison of the KEGG analysis of protein expression changes between AB1167 constructs exposed to microgravity **(Figure 2B),** analysis from AB1167_HSP10 constructs exposed to microgravity, **(Figure 6B)** and the 284 proteins which were differentially expressed by exposure to microgravity in the AB1167 constructs but not the HSP10 overexpressing constructs **(Figure 7)** highlights 3 pathways which were identified from the 284 proteins differentially expressed by exposure to microgravity in the AB1167 constructs but not the HSP10 overexpressing constructs (i.e. lysosome, other glycan degradation and glycosaminoglycan degradation) that are not present in **Figure 6B** but were effects identified in the initial comparisons of effects of microgravity on control constructs.

We infer that this represents a protective effect of HSP10 overexpression against deleterious effects of microgravity exposure acting in part through preservation of these 3 pathways. It is also apparent that HSP10 overexpression exerts these protective effects by modifying or priming the muscle in a way that partially mimics the effect of microgravity through changes in glycolytic capacity, structural proteins and mitochondrial proteins **(Figures 3 and 4).**

Thus, these data support the likelihood that HSP10 overexpression has modified the constructs sufficiently to adapt them to deal with the changes in energy and other requirements driven by microgravity exposure. Other data indicate that microgravity exposure leads to a decline in muscle mitochondrial capacity (16, 29, 30) and our data appear to indicate that HSP10 overexpression induces relevant changes in the mitochondrial proteome that modifies the microgravity-induced changes through mitigating stress responses, altering metabolic pathways and promoting mitochondrial gene expression and translation. Induction of heat shock proteins by pharmacological approaches has been proposed as an approach to mitigate age-related muscle loss (36) and the data presented here indicate that this approach may also be beneficial in reducing the deleterious adaptations to microgravity seen in skeletal muscle.

### Study Limitations

The samples analysed here represent a sub-group of the full MicroAge experiment undertaken on the ISS where the effects of electrical stimulation of contraction and of artificial gravity induced by centrifugation on proteomic profiles were also studied (17). Data from these additional manipulations of the 3D constructs (an additional 4 experimental groups) will be reported separately, but the bioinformatic analysis of all groups was undertaken simultaneously to ensure conformity of practice and application of appropriate statistical conditions on the whole study.

The need to remove proteins that were present in the hydrogel but shared sequences with human proteins found in the muscle constructs for the bioinformatic analysis, together with removal of proteins that were missing from one sample or were not present in at least 2 samples within a replicate group meant that the total number of proteins analysed was 5067 compared with the 7536 originally detected by mass spectrometry. Of the samples removed, 975 were removed because of being missing in one or more of the overall 8 experimental groups. In order to determine whether this may have biased the data by exclusion of proteins overrepresented within any specific biological processes, pathway enrichment analysis of the 975 proteins was undertaken (data not shown in detail). Some specific muscle proteins were found to be removed by this process such as key proteins involved in muscular contraction, but there was no over-emphasis in proteins from any particular pathways, and the pathway analysis did not identify any enriched pathways in the group of missing proteins suggesting no bias in interpretation was introduced by this process.

## Conflicts of Interest

The authors declare that the research was conducted in the absence of any commercial or financial relationships that could be construed as a potential conflict of interest.

## Author Contributions

Author SWJ prepared the article. Authors SWJ, SS and KFH generated the samples for LC-MS analysis. The Centre for Proteome Research (authors KA, PB and CE) performed the LC-MS analysis. The Computational Biology Facility (authors MH and AJ) performed the bioinformatic analysis of the LC-MS data and prepared the figures. Authors GN, WB, GO, KH and CM were responsible for hardware development activities. Authors KH, CM, and JH contributed equally in their critical appraisal of the manuscript. Authors MJJ and AM conceptualised the experiment, obtained the funding and edited the manuscript. All authors approved the final submitted article.

## Acknowledgements

This work was generously supported by the UK Space Agency through a grant from the UK Science and Technology facilities Council (grant number ST/S003061/1). The funder was not involved in the study design, data collection, interpretation, writing or the decision to submit the present article for publication. The authors acknowledge Dr Emily Johnson (Computational Biology Facility, University of Liverpool) for assistance with final manuscript preparation.

## Supplementary data

**Supplementary Figure S1. (i).** Lentiviral plasmid expression map for the overexpression of HSP10. The plasmid (HSP10-T2A-Bsd) was constructed with a T2A linker to achieve polycistronic expression of an EGFP reporter gene and the human HSP10 (HSPE1) target sequence. **(ii).** Optimising the stable overexpression of Heat Shock Protein 10 (HSP10) in human immortalised myoblasts (AB1167_CTL) by lentiviral transduction. (A-C) Percentage transduction efficiency was calculated by counting the number of cells that expressed the fluorescent EGFP reporter as proportion of the total number of cells within a field of view (FOV), a minimum of three separate FOVs were analysed per condition. (D) Representative (n of 3) brightfield and fluorescent (excitation 488nm, EGFP reporter) images of the AB1167_HSP10 myoblasts after 240 hours in blasticidin selection media (MOI 10-100), scale bar 100µm. (E) Representative western blot image demonstrating that HSP10 is overexpressed in the mutant cell line, as well showing the absence of EGFP in the control cell line (MOI 100). GAPDH was used as the loading control. (F) Gene expression data (RT-qPCR), illustrating increased expression of the HSP10 gene in the mutant cell line (MOI 100). All graphical values are expressed as a mean ± standard deviation (minimum n=3). Data were tested for Gaussian distribution using a Shapiro-Wilk test before statistical significance was determined using a student’s t-test with Welch’s correction. *p value < 0.05.

**Supplementary Figure S2**. “UpSet” plot showing the overlapping proteins in the different experimental groups and illustrating the extensive interactions in these sets of data.

**Supplementary Figure S3.** Heat maps of the top 20 MitoCarta 3.0 proteins modified by exposure of AB1167 constructs to microgravity (A) and exposure of AB11657_HSP10 constructs to microgravity (B).

**Supplementary Figure S4.** Heat maps of the top 20 MitoCarta 3.0 proteins modified by overexpression of HSP10 in AB1167 constructs in the GRE

**Supplementary Table 1.** Summary of sample protein concentration from each Experimental Unit (EU).

## Notes

https://github.com/CBFLivUni/MicroAge_Proteomics_paper

### Competing Interest Statement

The authors have declared no competing interest.

