## SupplementaryTable 1 for "MicroAge Mission: Effects of Microgravity and Heat Shock Protein 10 Overexpression on the Proteome of Human Tissue-Engineered Muscle Constructs - Implications for Skeletal Muscle Ageing"

| EU ID | Sample ID | µg or GRE | Cell Type | Protein Concentration (µg/µL) |
| --- | --- | --- | --- | --- |
| EU19 | 19.1 | µg | AB1167_CTL | 0.766 |
|  | 19.2 |  |  | 0.846 |
|  | 19.3 |  |  | 0.669 |
| EU20 | 20.1 | µg | AB1167_CTL | 0.5681 |
|  | 20.2 |  |  | 0.6124 |
|  | 20.3 |  |  | 1.047 |
| EU23 | 23.1 | µg | AB1167_HSP10 | 0.893 |
|  | 23.2 |  |  | 1.708 |
|  | 23.3 |  |  | 0.764 |
| EU 24 | 24.1 | µg | AB1167_HSP10 | 0.764 |
|  | 24.2 |  |  | 1.183 |
|  | 24.3 |  |  | 0.3519 |
| EU39 | 39.1 | GRE | AB1167_CTL | 0.845 |
|  | 39.2 |  |  | 0.739 |
|  | 39.3 |  |  | 0.744 |
| EU40 | 40.1 | GRE | AB1167_CTL | 0.2200 |
|  | 40.2 |  |  | 0.1029 |
|  | 40.3 |  |  | 0.2891 |
| EU43 | 43.1 | GRE | AB1167_CTL | 0.5126 |
|  | 43.2 |  |  | 0.956 |
|  | 43.3 |  |  | 0.5690 |
| EU44 | 44.1 | GRE | AB1167_CTL | 0.5060 |
|  | 44.2 |  |  | 1.041 |
|  | 44.3 |  |  | 0.5720 |
| EU47 | 47.1 | GRE | AB1167_HSP10 | 0.4772 |
|  | 47.2 |  |  | 0.785 |
|  | 47.3 |  |  | 0.2934 |
| EU48 | 48.1 | GRE | AB1167_HSP10 | 0.629 |
|  | 48.2 |  |  | 0.848 |
|  | 48.3 |  |  | 1.277 |

***Supplementary Table 1:*** *Summary of sample protein concentrations from each Experiment Unit (EU)*

Abbreviations: EU, Experiment Unit; µg, microgravity; GRE, Ground Reference Experiment.
