## Supplementary Figures for "MicroAge Mission: Effects of Microgravity and Heat Shock Protein 10 Overexpression on the Proteome of Human Tissue-Engineered Muscle Constructs - Implications for Skeletal Muscle Ageing"

### Slide 1
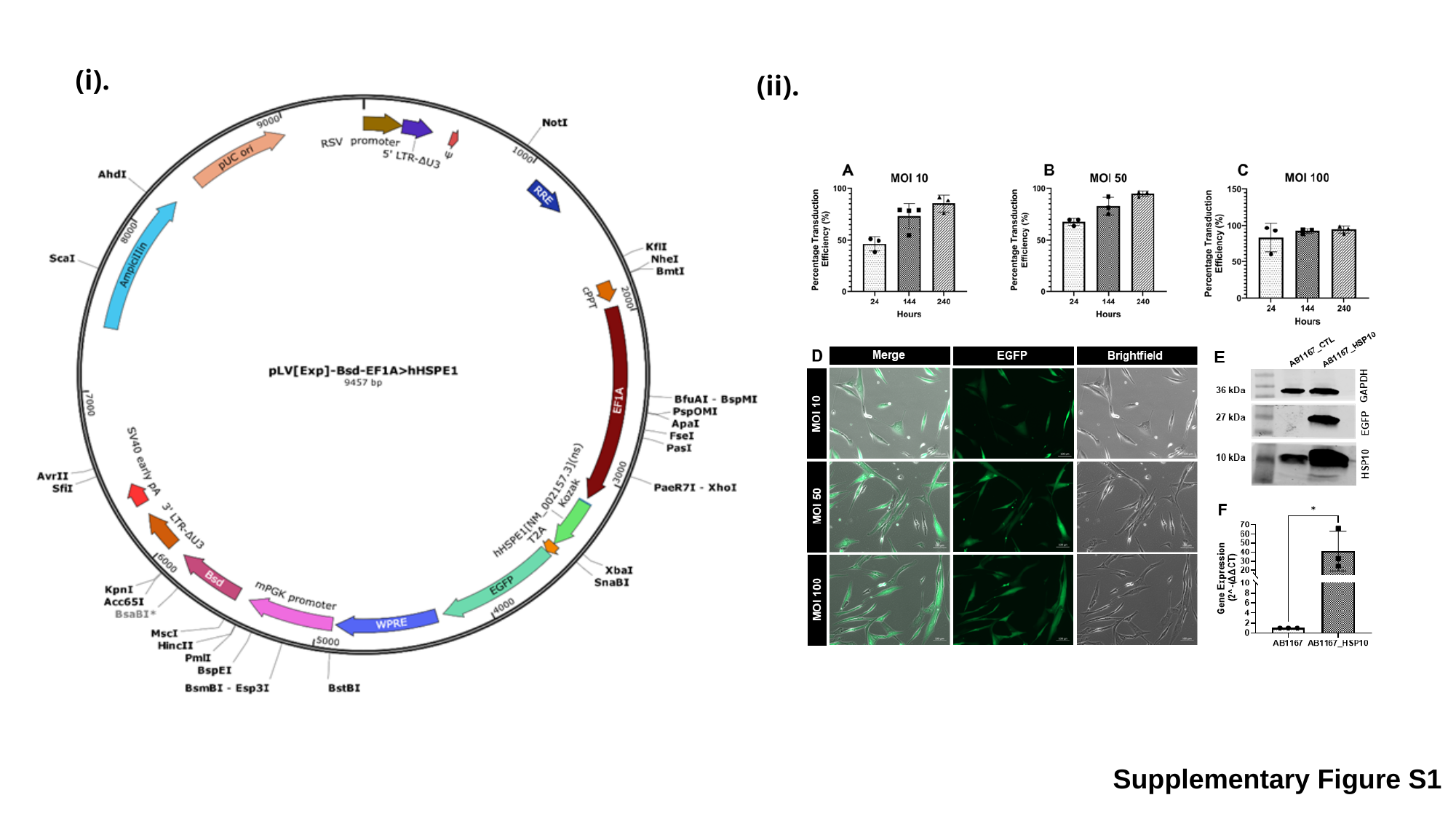

(i).
(ii).
Supplementary Figure S1

### Slide 2
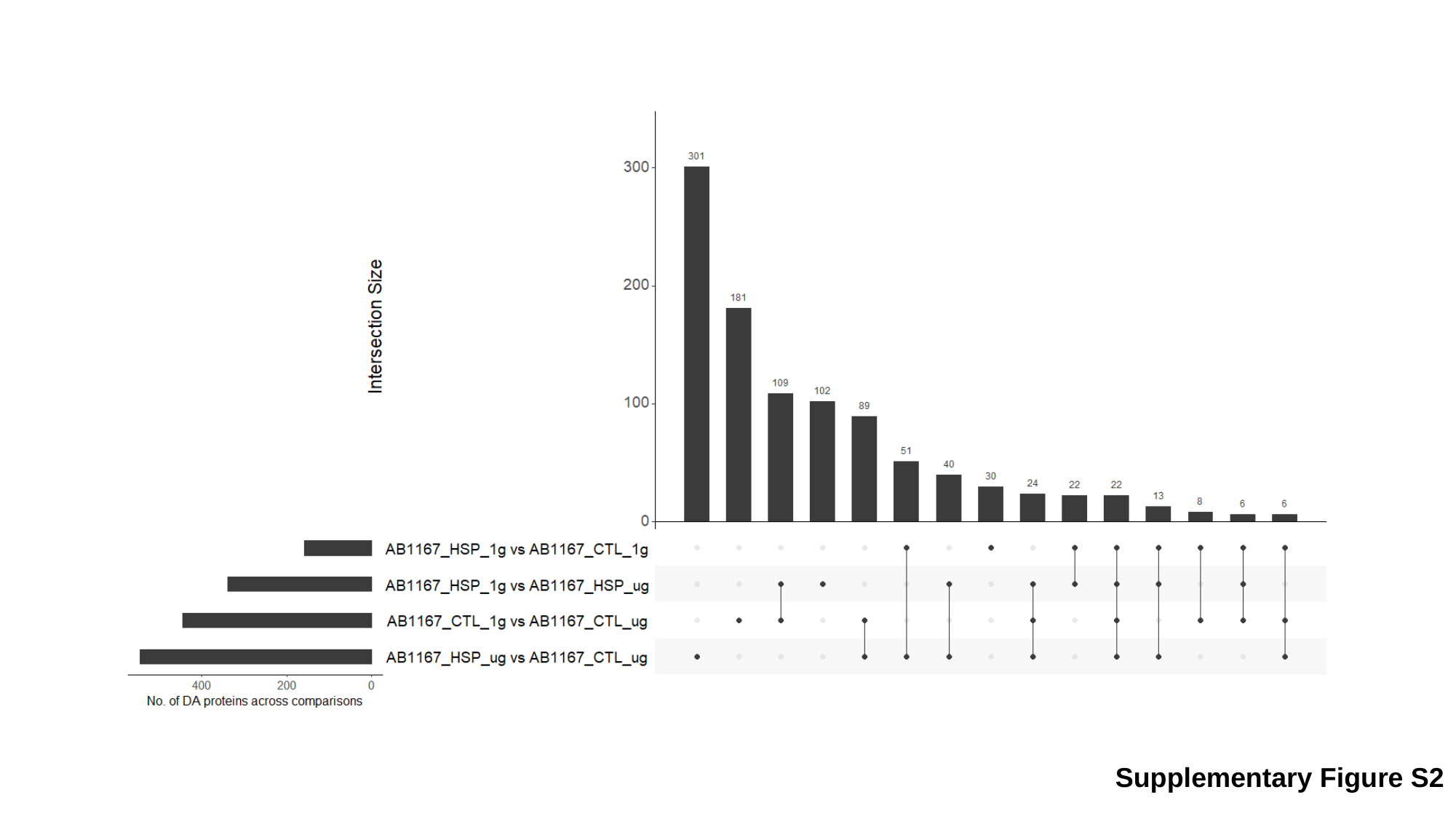

Supplementary Figure S2

### Slide 3
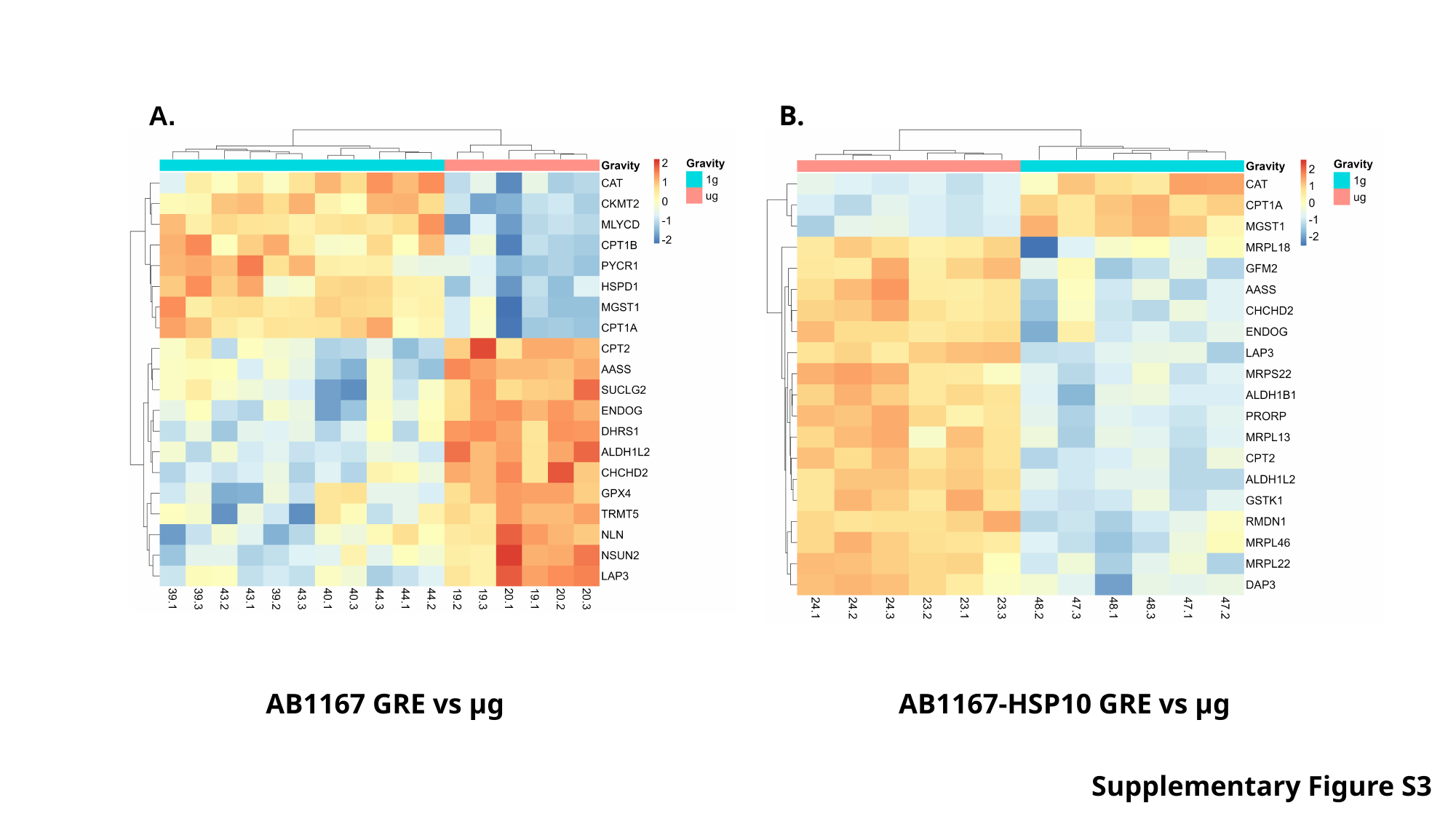

A.
B.
AB1167 GRE vs µg
AB1167-HSP10 GRE vs µg
Supplementary Figure S3

### Slide 4
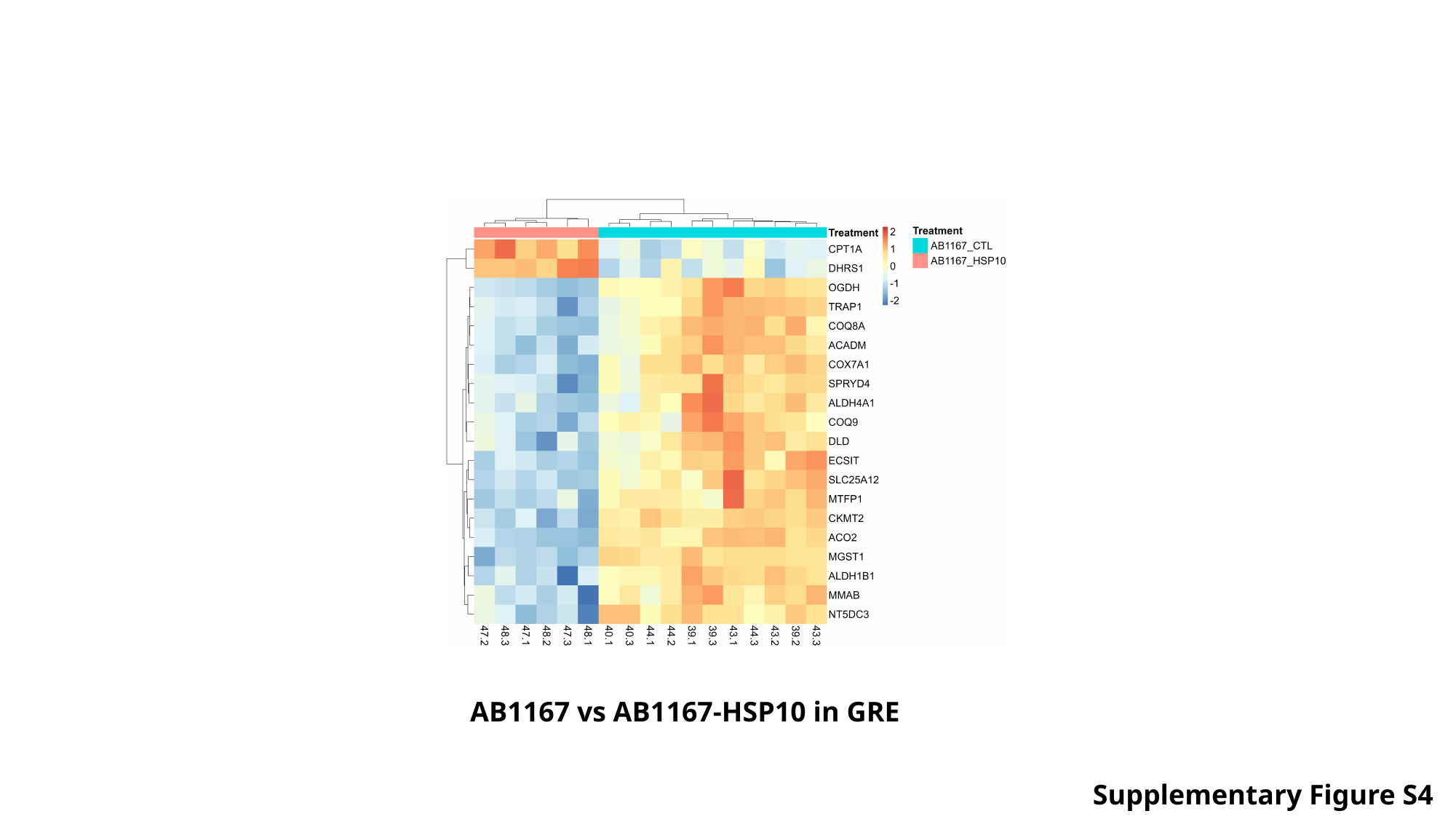

AB1167 vs AB1167-HSP10 in GRE
Supplementary Figure S4
